# Establishment of a stable SARS-CoV-2 replicon system for application in high-throughput screening

**DOI:** 10.1101/2021.12.23.474055

**Authors:** Tomohisa Tanaka, Akatsuki Saito, Tatsuya Suzuki, Yoichi Miyamoto, Kazuo Takayama, Toru Okamoto, Kohji Moriishi

**Author notes:** **Corresponding author:** Kohji Moriishi, D.V.M., Ph.D., Department of Microbiology, Faculty of Medicine, Graduate Faculty of Interdisciplinary Research, University of Yamanashi, 1110 Shimokato, Chuo-shi, Yamanashi 409-3898, Japan.

## Abstract

Experiments with severe acute respiratory syndrome coronavirus 2 (SARS-CoV-2) are limited by the need for biosafety level 3 (BSL3) conditions. A SARS-CoV-2 replicon system rather than an *in vitro* infection system is suitable for antiviral screening since it can be handled under BSL2 conditions and does not produce infectious particles. However, the reported replicon systems are cumbersome because of the need for transient transfection in each assay. In this study, we constructed a bacterial artificial chromosome vector (the replicon-BAC vector) including the SARS-CoV-2 replicon and a fusion gene encoding *Renilla* luciferase and neomycin phosphotransferase II, examined the antiviral effects of several known compounds, and then established a cell line stably harboring the replicon-BAC vector. Several cell lines transiently transfected with the replicon-BAC vector produced subgenomic replicon RNAs (sgRNAs) and viral proteins, and exhibited luciferase activity. In the transient replicon system, treatment with remdesivir or interferon-β but not with camostat or favipiravir suppressed the production of viral agents and luciferase, indicating that luciferase activity corresponds to viral replication. VeroE6/Rep3, a stable replicon cell line based on VeroE6 cells, was successfully established and continuously produced viral proteins, sgRNAs and luciferase, and their production was suppressed by treatment with remdesivir or interferon-β. Molnupiravir, a novel coronavirus RdRp inhibitor, inhibited viral replication more potently in VeroE6/Rep3 cells than in VeroE6-based transient replicon cells. In summary, our stable replicon system will be a powerful tool for the identification of SARS-CoV-2 antivirals through high-throughput screening.

## Introduction

Coronavirus disease 2019 (COVID-19), which is caused by severe acute respiratory syndrome coronavirus 2 (SARS-CoV-2), poses a global threat to public health. Up to 80% of people infected with SARS-CoV-2 exhibit asymptomatic cases or mild-to-moderate respiratory illness, while some patients progress to severe pneumonia, acute respiratory distress, septic shock and multiple organ failure (Cao, 2020; Huang et al., 2020). As of November 2021, more than 200 million people have been affected by COVID-19, and approximately 5 million deaths have been reported worldwide (Yang and Rao, 2021). Some of the current vaccines against the SARS-CoV-2 spike protein are highly effective, while several amino acid residues in the viral spike protein have been reported to be mutated, leading to impairment of vaccine efficacy (Collier et al., 2021; Harvey et al., 2021). Neutralizing antibody cocktails targeting the spike protein and small chemicals such as remdesivir, molnupiravir and some protease inhibitors have been developed for the treatment of COVID-19 (Bakowski et al., 2021; Choy et al., 2020; Group, 2020; Sheahan et al., 2020). However, more effective antivirals and vaccines are still needed.

SARS-CoV-2 is an enveloped, positive-sense, single-stranded RNA virus belonging to the genus *Betacoronavirus*, subgenus *Sarbecovirus*. Two viral polyproteins (pp1a and 1ab) are cleaved into 16 nonstructural proteins, which constitute the replication and transcription complex to initiate viral RNA replication and transcription (Snijder et al., 2016; Yang and Rao, 2021). Nsp12 possesses RNA polymerase activity and forms a tripartite complex with nsp7 and nsp8 to produce the RNA-dependent RNA polymerase (RdRp) complex (Snijder et al., 2016). Nucleocapsid (N) protein binds to viral genomic RNA to form the ribonucleocapsid and helps regulate discontinuous transcription of subgenomic RNAs (sgRNAs) through RNA chaperone activity (Grossoehme et al., 2009; McBride et al., 2014; Zuniga et al., 2010).

Since experimental infection with SARS-CoV-2 must be restricted under the condition of biosafety level 3 (BSL3), BSL2-compatible SARS-CoV-2 replicon systems have been developed instead of *in vitro* infection models for basic research and antiviral screening (He et al., 2021; Kotaki et al., 2021; Nguyen et al., 2021; Zhang et al., 2021). Nguyen et al. prepared a bacterial artificial chromosome (BAC) vector encoding SARS-CoV-2 replicon RNA under the control of the CMV promoter (Nguyen et al., 2021), while others have made transient replicon systems using *in vitro*-transcribed SARS-CoV-2 replicon RNA (He et al., 2021; Kotaki et al., 2021; Zhang et al., 2021). In each replicon system, the expression of luciferase or a fluorescent reporter gene of replicon

RNA is induced in a replication-dependent manner. These replicon systems could be applicable to screening for antiviral compounds targeting SARS-CoV-2 replication, although transient transfection with these vectors or RNAs is still needed for each replicon assay, leading to cumbersome application for studies such as high-throughput screening. Thus, establishment of a cell line harboring stable replication of the SARS-CoV-2 replicon is required for drug screening.

In this study, we prepared a BAC vector encoding SARS-CoV-2 reporter replicon RNA with a fusion gene encoding *Renilla* luciferase and neomycin phosphotransferase II (the replicon-BAC vector), examined the antiviral effects of remdesivir, molnupiravir, type I interferon (IFN), camostat and favipiravir on viral replication in the transient system using the replicon-BAC vector, and established a cell line that stably supports the replication of the replicon. Furthermore, we examined the production of viral factors and the antiviral effect of the reported compounds in the stable replicon cell line. Our stable replicon system for SARS-CoV-2 will be a powerful tool for high-throughput screening for clinically available antivirals.

## Results

### Development of the BAC vector encoding the SARS-CoV-2 reporter replicon

Several groups have reported the development of SARS-CoV-2 replicon systems (Kotaki et al., 2021), while the development of a cell clone stably replicating the replicon has not yet been reported. In this study, we constructed a SARS-CoV-2-based selectable reporter replicon encoding a fusion gene encoding *Renilla* luciferase and neomycin phosphotransferase (Reo) to establish a stable SARS-CoV-2 replicon cell line. The reporter replicon cDNA was engineered with a BAC due to the large size and to achieve stability of the coronavirus replicase gene (ORF1a, 1b), according to reports described by others (Almazan et al., 2006; Nguyen et al., 2021). We employed the circular polymerase extension reaction (CPER) method to assemble the BAC vector containing SARS-CoV-2 replicon cDNA (Edmonds et al., 2013; Torii et al., 2021). Eight fragments covering the region encoding nonstructural polyproteins were subcloned into pCR-blunt (Invitrogen) as templates for fragments F1 to F8 (Fig. 1A). Fragment F9, which is composed of, in order, a spacer (AscI site), the 3’-end portion of ORF8, transcription-regulating sequence (TRS)-body (TRS-B), the *Reo* gene, a spacer (BamHI site), the 3’-end portion of ORF8, and TRS-B, was subcloned into pCR-blunt (Fig. 1A, S1A). Fragments F1 to F10, each of which possesses approximately 30-nt overlapping ends for two neighboring fragments, were amplified by PCR using individual primers (Table 1) and subjected to CPER. The resulting BAC vector was designated pBAC-SCoV2-Rep-Reo (Fig. 1A).

**Figure 1.**
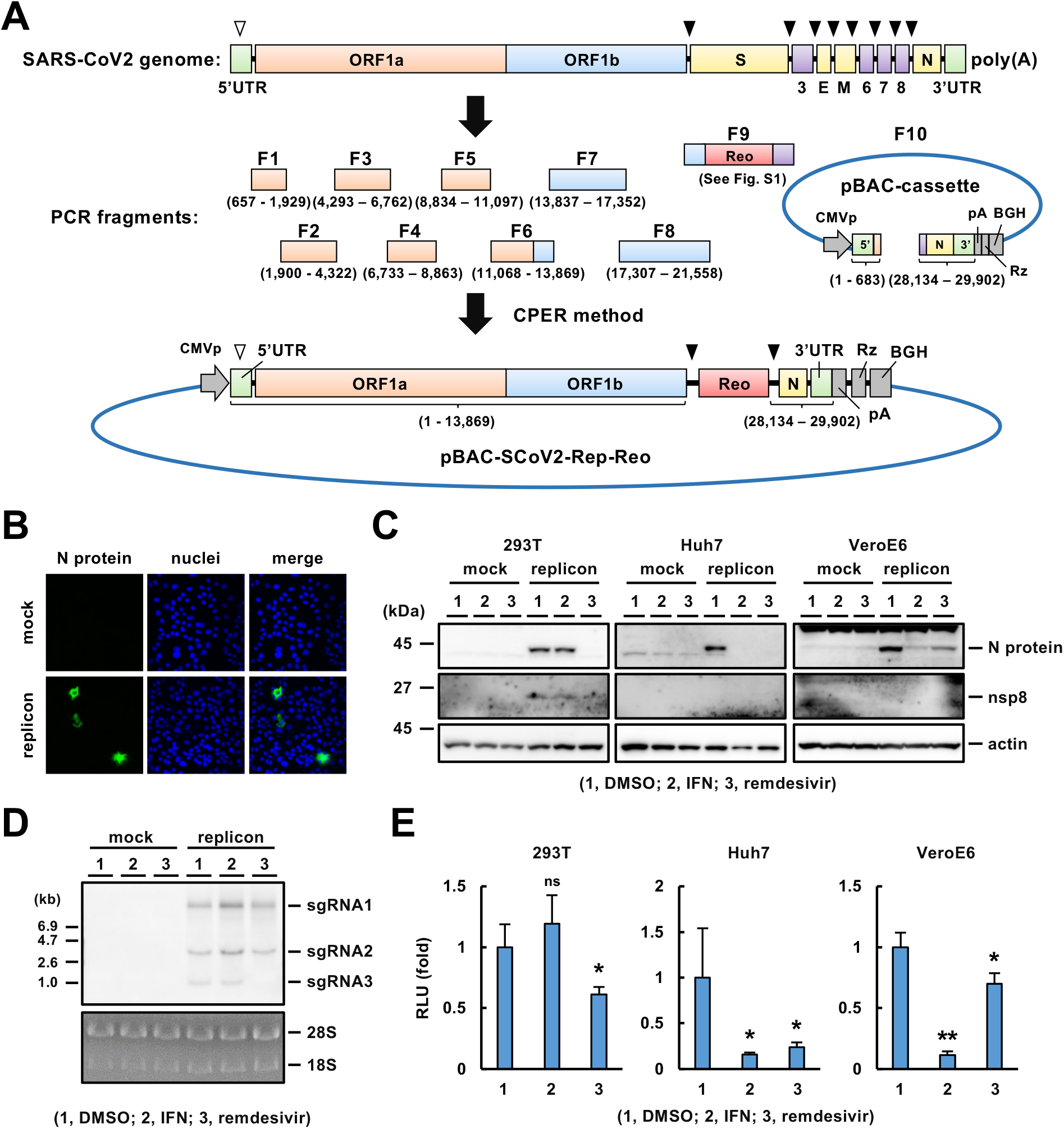
Construction of the BAC vector encoding the SARS-CoV-2 reporter replicon. (A) A schematic diagram of the strategy for cloning the BAC vector encoding the SARS-CoV-2 reporter replicon. PCR fragments F1 to F10 were amplified from subcloning or cassette vectors by PCR using the primers listed in Table 1 as described in Materials and Methods and then subjected to CPER. The resulting vector was designated pBAC-SCoV2-Rep-Reo. The numbers inside the parentheses represent the nucleotide positions in the SARS-CoV-2/Hu/DP/Kng/19-020 genome. TRSs in the leader sequence of the 5’ UTR (TRS-L) and in the genome body (TRS-B) are indicated as a white arrowhead and a black arrowhead, respectively. The blue line indicates the pSMART-BAC v2.0 vector sequence. Reo, a fusion gene consisting of *Renilla* luciferase and neomycin phosphotransferase; CMVp, cytomegalovirus promoter; 5’, 5’ UTR; 3’, 3’ UTR; pA, synthetic 25-poly(A); Rz, hepatitis delta virus ribozyme; BGH, bovine growth hormone termination and polyadenylation sequence. Refer to Fig. S1 for detailed information on fragment F9 and the intermediate sequence between the ORF1ab and N genes in pBAC-SCoV2-Rep-Reo. (B) Immunostaining of N protein in VeroE6 cells transfected with pBAC-SCoV2-Rep-Reo or mock. These cells were fixed at 24 h post-transfection. Nuclei were stained with DAPI. (C) Western blotting of viral proteins. VeroE6, Huh7 and 293T cells were transfected with the replicon-BAC vector or mock, treated with DMSO (0.1%), IFN-β (100 U/ml) or remdesivir (0.5 μM for 293T; 5 μM for Huh7 and VeroE6) at 6 h post-transfection, and harvested at 54 h post-transfection. The cell lysates were subjected to Western blotting using specific antibodies against the N protein, nsp8 or actin. (D) Northern blotting of sgRNAs. The replicon-BAC vector was transfected into 293T cells as described in (C). The resulting cells were treated with each reagent and subjected to Northern blotting. Ribosomal RNAs were stained with EtBr. (E) Luciferase activities in the cells transfected with the replicon vector or mock. The replicon-BAC vector was transfected into 293T cells as described in (C). The resulting cells were treated with each reagent and subjected to a luciferase assay. Statistical significance was calculated compared to the mock group (*,*p* < 0.05; **,*p* < 0.01; ns, no significance).

**Table 1.**
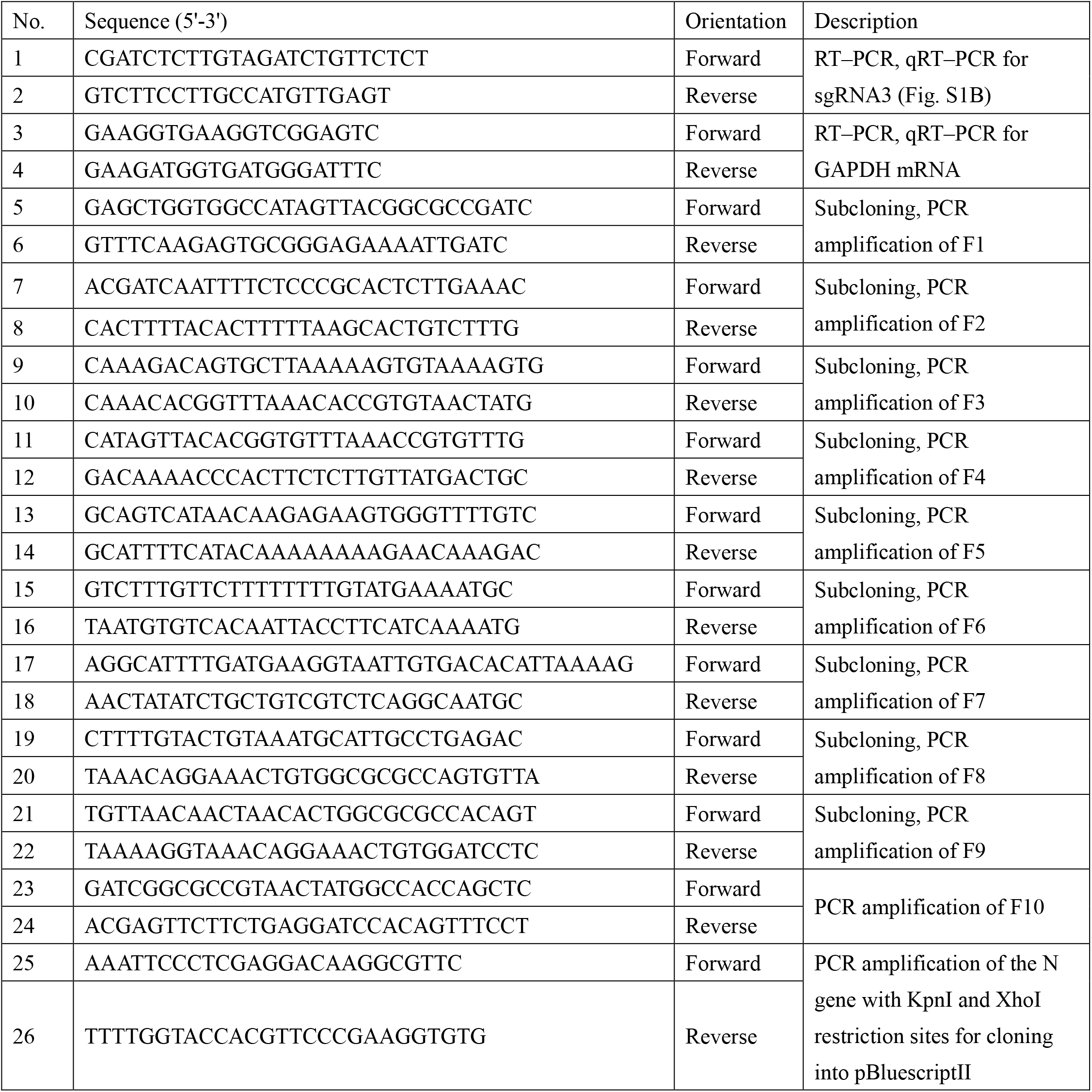
Primer sequences.

### Selection of the replication-competent pBAC-SCoV2-Rep-Reo

The BAC vector-encoded replicon cDNA is transcribed from the CMVp by the host RNA polymerase complex into the replicon mRNA, which is processed by ribozyme-mediated self-cleavage to generate full-length replicon RNA (Fig. S1B). The full replicon RNA contains three TRSs: one TRS in the leader sequence of the 5’ untranslated region (UTR) (TRS-L) and two TRSs upstream of the Reo and N genes (TRS-B). TRSs allow recombination between TRS-L and TRS-B to produce sgRNAs in a replication-dependent manner (Fig. S1B). For the selection of transformant colonies carrying the CPER product (Fig. S1C), each DNA preparation extracted from 20 colonies was introduced into 293T cells by lipofection. Following transfection, RT-PCR specific for sgRNA3 using primers 1 and 2 (Fig. S1B, Table 1) and luciferase assays were carried out to find replication-competent BAC clones. Ten clones were positive for both sgRNA3 transcription and Reo translation (Fig. S1C). We purified the DNA preparation from clone number 3, confirmed cloning junctions in CPER and employed this preparation for subsequent experiments.

### Transient expression of the SARS-CoV-2 reporter replicon in cell culture

We examined whether pBAC-SCoV2-Rep-Reo was active in a transient replicon assay. N protein was detected in VeroE6 cells transfected with the replicon-BAC vector (Fig. 1B). We transiently transfected the replicon-BAC vector into 293T, Huh7 and VeroE6 cells and then cultured the resulting cells in the presence of IFN-β or remdesivir (Lei et al., 2020; Zhao et al., 2021a). N protein was expressed in the cell lines transfected with the replicon-BAC vector (Fig. 1C), and its expression level was reduced by treatment with IFN-β or remdesivir, depending on the cell line (Fig. 1C), suggesting replication-dependent expression of the N protein. Treatment with IFN-β reduced N protein expression in Huh7 and VeroE6 cells but not 293T cells. In contrast, treatment with remdesivir reduced N protein expression in 293T and Huh7 cells but weakly in VeroE6 cells (Fig. 1C). Additionally, nsp8, a component of the viral replicase complex, was detected in the transfected 293T cells, and its expression was reduced by remdesivir, suggesting that polyproteins 1a and 1ab were expressed and processed into mature nonstructural proteins (Fig. 1C). Nsp8 was undetectable in Huh7 and VeroE6 cells by Western blotting, probably due to limited transfection efficiency in these cell lines (Fig. 1C). Additionally, we analyzed the replicon RNA extracted from the transfected 293T cells by Northern blotting using a probe specific for the N gene. Three sgRNAs (sgRNA1, 2 and 3, Fig. S1B) with distinct predicted sizes were detected by Northern blotting (Fig. 1D). Interestingly, the sgRNA3 level was clearly decreased by treatment with remdesivir but not by treatment with IFN-β in 293T cells (Fig. 1D). The reduction in the sgRNA3 level by remdesivir treatment was confirmed quantitatively by qRT-PCR (Fig. S1E). The effects of IFN-β and remdesivir on the level of sgRNA3 were well correlated with their effects on the level of N protein in 293T, Huh7 and VeroE6 cells (Fig. 1C, Fig. S1E). Luciferase activities in 293T, Huh7 and VeroE6 cells transfected with the replicon-BAC vector were significantly reduced by treatment with IFN-β and/or remdesivir (Fig. 1E). Consistent with the Western blotting and qRT-PCR data (Fig. 1C, Fig. S1E), IFN-β and remdesivir significantly impaired luciferase expression in the Huh7 and VeroE6 cell lines transfected with the replicon-BAC vector, while viral replication in 293T cells was resistant to IFN-β (Fig. 1E). These data suggest that SARS-CoV-2 replication can be monitored by the reporter assay.

Recently, camostat mesylate, an inhibitor of TMPRSS2, was shown to inhibit SARS-CoV-2 entry (Cheng et al., 2020; Hoffmann et al., 2021; Hoffmann et al., 2020). Favipiravir (Avigan), a nucleoside analog, has been shown to have antiviral activity in *in vivo* assays and clinical studies (Kaptein et al., 2020; Martinez, 2020) and to suppress viral replication via inhibition of viral RdRp at a high dose up to 100 μM (Shannon et al., 2020; Zhao et al., 2021a), whereas some reports show no obvious inhibition of virus proliferation upon treatment with favipiravir (Choy et al., 2020; Pizzorno et al., 2020). Luciferase activity was not reduced by treatment with 10 μM favipiravir in 293T cells transfected with the replicon-BAC vector (Fig. S1F), suggesting that the transient replicon system is applicable to the assessment of antiviral effects on specifically SARS-CoV-2 replication.

### Establishment of a VeroE6-based cell line harboring a functional SARS-CoV-2 replicon with luciferase expression

We attempted to establish a stable replicon cell line using pBAC-SCoV2-Rep-Reo. Because the replicon-BAC vector was designed to confer neomycin resistance to cells, we cultured Huh7 and VeroE6 cells transfected with the replicon vector in medium containing G418. A number of colonies were generated by the transfection of VeroE6 cells with the replicon-BAC vector (Fig. S2A) but not by transfection of Huh7 cells with the vector (data not shown). We established 14 cell clones from these colonies. Since the expression of the reporter and sgRNAs requires TRS-mediated discontinuous transcription, we evaluated cellular luciferase activity in each clone. A higher level of luciferase activity was observed in clone nos. 1, 3, 4, 6 and 13 than in the other clones (Fig. S2B; nos. 1, 3, 4, 6 and 13). However, sgRNA3 was detected in clone no. 3 but not in the other clones (Fig. S2C). Therefore, clone no. 3 was analyzed further as a stable replicon cell line, designated VeroE6/Rep3. To assess the sustainability of persistent reporter expression in VeroE6/Rep3 cells, we serially passaged the cells in the presence or absence of G418. The luciferase activity in VeroE6/Rep3 cells was maintained for at least 4 weeks in the presence of G418 (Fig. S2D). The luciferase activity was also maintained but decayed when the cells were cultured for 2 weeks in the absence of G418 (Fig. S2D). Therefore, it is better to maintain VeroE6/Rep3 cells in the presence of G418, while the medium can also be replaced with G418-free medium during short-term assays.

To determine whether the replicon was actively replicated in the VeroE6/Rep3 cell line, we evaluated the intracellular levels of N protein and sgRNAs in the presence or absence of IFN-β or remdesivir. The expression of N protein in VeroE6/Rep3 cells was at a level similar to that in VeroE6 cells transiently transfected with the vector (Fig. 2A) and was reduced by treatment with IFN-β or remdesivir (Fig. 2A), suggesting that VeroE6/Rep3 cells express N protein in a replication-dependent manner. Northern blotting data indicated that sgRNAs 1, 2 and 3 were expressed in the VeroE6/Rep3 cell line but not in the parent VeroE6 cell line (Fig. 2B). N protein was expressed in VerpE6/Rep3 cells, but not in VeroE6 cells (Fig. 2C). Treatment with IFN-β decreased the levels of all sgRNAs, while treatment with remdesivir decreased the amount of sgRNA3 more potently than those of sgRNAs 1 and 2 (Fig. 2D). Furthermore, treatment with IFN-β or remdesivir reduced luciferase expression in VeroE6/Rep3 cells, although the inhibitory effect of remdesivir was lower than that of IFN-β (Fig. 2E), which is consistent with their effects on the level of N protein (Fig. 2A). These data suggest that luciferase activity is correlated with the replication level in VeroE6/Rep3 cells.

**Figure 2.**
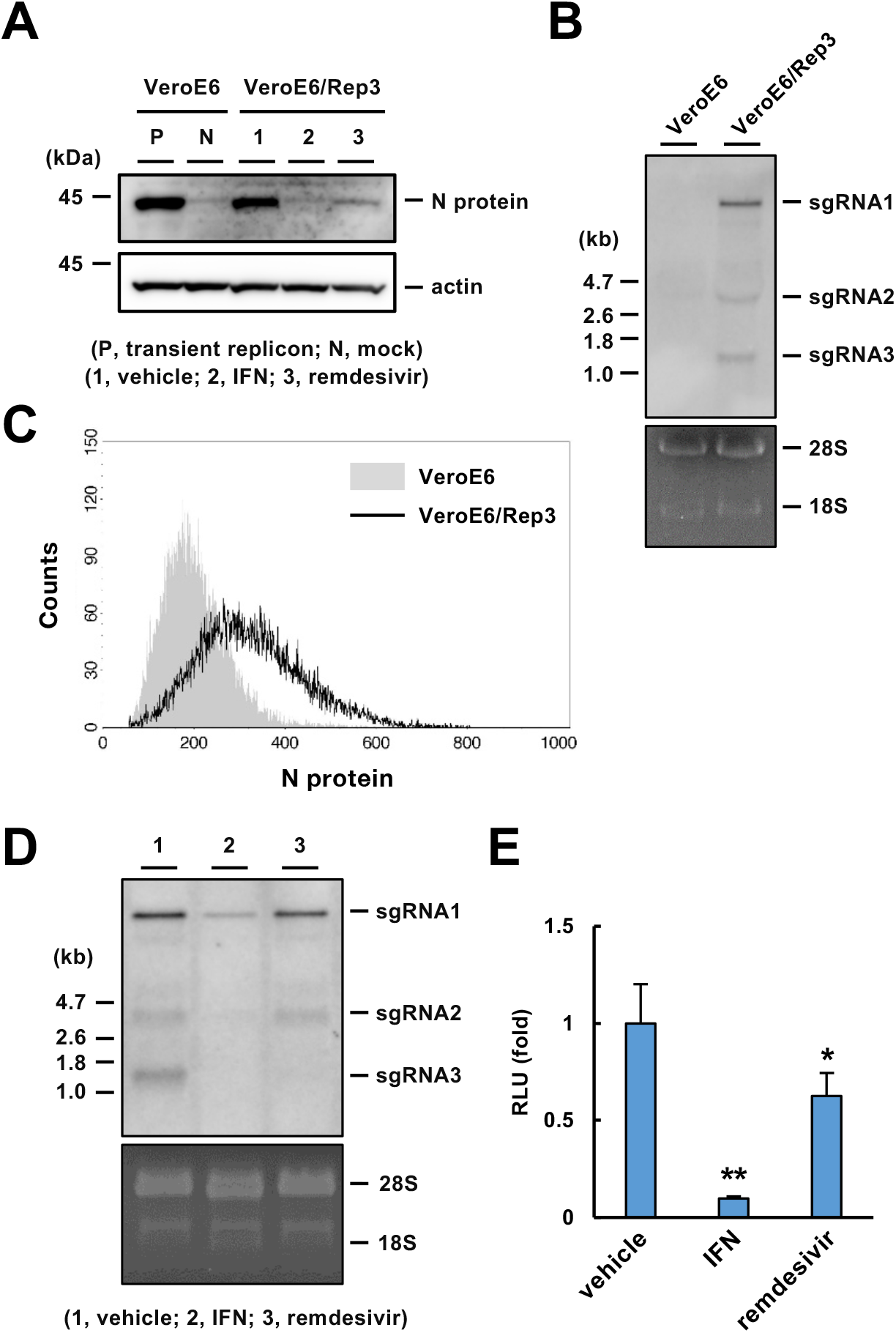
Establishment of the VeroE6/Rep3 stable replicon cell line. (A) Western blotting of viral proteins in the VeroE6/Rep3 cell line. The VeroE6/Rep3 cell line was treated with DMSO (0.1%), IFN-β (100 U/ml) or remdesivir (5 μM), and the cells were harvested 48 h after treatment. The cell lysates were subjected to Western blotting. Lysates of VeroE6 cells transfected with pBAC-SCoV2-Rep-Reo (P) or mock (N) were prepared as controls. (B) Northern blotting of sgRNAs. Total RNA was extracted from VeroE6 and VeroE6/Rep3 cells and then subjected to Northern blotting using an RNA probe targeting the N-coding region. Ribosomal RNAs were stained with EtBr. (C) FACS analysis of N protein expression in the VeroE6/Rep3 cell line. (D) Northern blotting using an RNA probe for the SARS-CoV-2 N gene. Total RNA was extracted from the VeroE6/Rep3 cell line that was treated with DMSO (0.1%), IFN-β (100 U/ml) or remdesivir (5 μM) for 48 h. Ribosomal RNA was stained with EtBr. (E) Effect of antivirals on viral replication determined by evaluation of luciferase activity in the VeroE6/Rep3 cell line. VeroE6/Rep3 cells were treated with DMSO (0.1%), IFN-β (100 U/ml) or remdesivir (5 μM) and then harvested 48 h after treatment. The cell lysates were subjected to a luciferase assay. Statistical significance was determined compared to the mock group (*, p < 0.05; **, p < 0.01).

### Effect of EIDD-2801 on SARS-CoV-2 replication

EIDD-2801 (molnupiravir), a prodrug of the ribonucleoside analog β-D-N4-hydroxycytidine (NHC), was recently reported to exhibit potent antiviral activity against SARS-CoV-2 *in vivo* and *in vitro* (Bakowski et al., 2021; Painter et al., 2021; Sheahan et al., 2020; Wahl et al., 2021). The *in vitro* reporter system for the analysis of SARS-CoV-2 RdRp activity showed that EIDD-2801 effectively inhibited RdRp activity with an IC_50_ value of 0.22 μM (Zhao et al., 2021a). We examined the effect of EIDD-2801 on viral replication using our transient and stable replicon systems. The luciferase activity was dose-dependently reduced by treatment with EIDD-2801 in Huh7 cells but not 293T cells transiently transfected with the BAC vector (Fig. 3A). EIDD-2801 effectively inhibited replication in Huh7 cells but not remarkably in VeroE6 cells (Fig. 3A). The IC_50_ value of EIDD-2801 in transfected Huh7 cells was calculated to be approximately 2.2 μM. On the other hand, the luciferase activity was also reduced by EIDD-2801 treatment in a dose-dependent manner in VeroE6/Rep3 cells (Fig. 3B), and the drug effect appeared more remarkable in VeroE6/Rep3 cells than in VeroE6 cells transfected with the replicon-BAC vector (Fig. 3A, B). The IC_50_ value of EIDD-2801 in VeroE6/Rep3 cells was calculated to be approximately 4.3 μM. These data suggest that our transient and stable replicon systems provide insight into the replication mechanisms of SARS-CoV-2 and contribute to the development of antiviral agents against SARS-CoV-2 under BSL-2 conditions.

**Figure 3.**
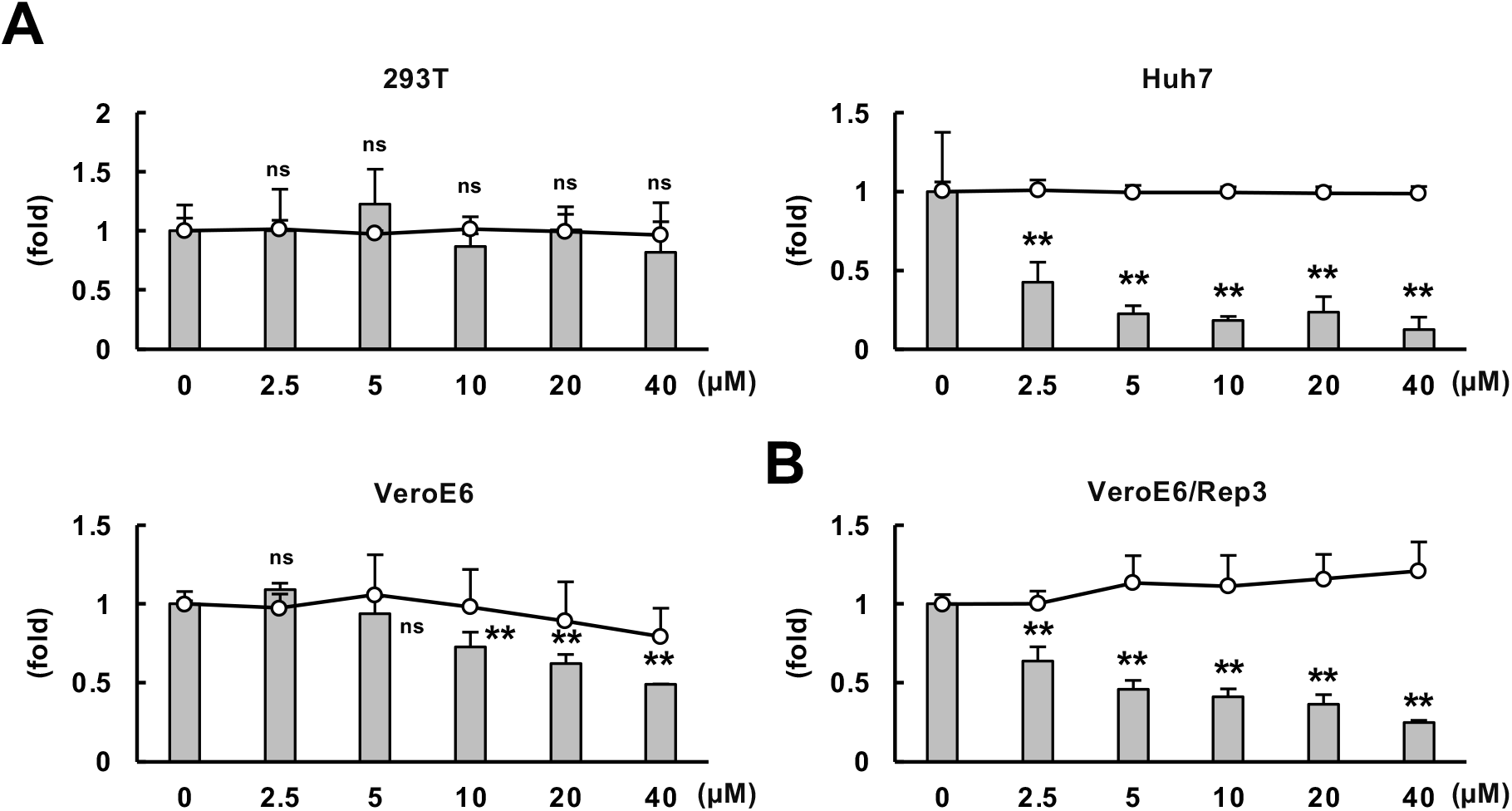
The effect of EIDD-2801 on replication evaluated in the transient and stable replicon models. (A) Analysis of the antiviral effect of EIDD-2801 by transient replicon assays. Huh7, 293T and VeroE6 cells were transfected with the replicon-BAC vector and treated with the indicated concentration of EIDD-2801 at 6 h post-transfection. The resulting cells were subjected to a luciferase or an MTS assay at 54 h post-transfection. The luciferase activities (columns) and cell viabilities (lines) were measured and are shown as the values relative to those of the vehicle-treated group. (B) Analysis of the antiviral effect of EIDD-2801 by stable replicon assay. VeroE6/Rep3 cells were incubated in the presence of the indicated concentrations of EIDD-2801 for 48 h. The luciferase activities (columns) and cell viabilities (lines) were measured and are shown as the values relative to those of the vehicle-treated group. (A, B) Statistical significance was calculated compared to the mock group (*, p < 0.05; **, p < 0.01; ns, no significance). There was no significant difference in cell viability (not shown).

## Discussion

One of the advantages of the CPER method is the ease of modification; site-directed mutation in infectious or replicon cDNA of SARS-CoV-2 can be achieved through the modification of individual subcloning vectors. Several therapeutic agents for SARS-CoV-2 are eagerly anticipated and will be put into practical use, while antiviral administration potentially involves the risk of the development of drug-resistant variants. The construction of replicon vectors by the CPER method provides an immediate response to the investigation of drug resistance and other novel mutants. We showed that the inhibitory effects of remdesivir, IFN-β and EIDD-2801 on replication varied among 293T, Huh7 and VeroE6 cells (Fig. 1C, E, Fig. 3). The differences in drug efficacy may be attributed to the cell-type characteristics of membrane permeability, efflux, metabolic pathway activation and IFN-β response. Thus, the option of cell types will be advantageous for the exploration of viral replication and drugs. It would be burdensome to conduct a DNA transfection step for the transient replicon system and to prepare a sufficient amount of high-grade BAC vector because a BAC vector with a large molecular size is expected to be prepared at a low yield. On the other hand, stable replicon systems, such as the VeroE6/Rep3 cell line, do not require transfection for each assay or vector preparation. Thus, the stable replicon system seems to be appropriate for high-throughput screening rather than the transient replicon system. The dose dependency of the reduction in the replicon replication level was clearer in VeroE6/Rep3 cells than in VeroE6 cells harboring the transient replicon (Fig. 3). In the transient replicon system, heterogeneity in transfection efficiency and cytotoxicity of transfection may lead to variability in the data. It will be important to choose between the two models depending on the purpose of the experiment.

Unfortunately, we could not establish another cell line persistently expressing the replicon in this study. The replicon vector was unstable in the Huh7 cell line and eradicated during a cloning step. Furthermore, a recent report suggested that the establishment of stable replicon cells is unsuccessful due to the toxicity derived from a viral protein or a replication step (He et al., 2021). Among the viral proteins translated from replicon sgRNAs, nonstructural protein 1 (nsp1) has been shown to be a viral cytotoxic factor (Yuan et al., 2020). Nsp1 plays an important role in the shut-off of host protein synthesis during SARS-CoV-2 replication (Thoms et al., 2020). Additionally, its expression has been shown to trigger cell cycle arrest and cell death (Shen et al., 2021; Thoms et al., 2020). Thus, nsp1-induced cytotoxicity in replicon-expressing cells is thought to interfere with the establishment of stable replicon cells. On the other hand, VeroE6 cells are known to lack type I IFN production and the expression of key factors in cell cycle regulation, including cyclin-dependent kinase inhibitors 2A and 2B (Osada et al., 2014). The characteristics of VeroE6 cells may contribute to the establishment of a stable cell line harboring the SARS-CoV-2 replicon.

The data in Figs. 1D and 2D indicated that treatment with remdesivir potently reduced the synthesis of sgRNA3 more than that of other sgRNAs in the transient and stable replicon models. Remdesivir is a nucleoside analog prodrug that is converted into the active triphosphate form (Siegel et al., 2017). The triphosphate form is used by viral RdRp instead of nucleoside triphosphates, resulting in subsequent RdRp stalling (Gordon et al., 2020; Kokic et al., 2021). Furthermore, remdesivir has been shown to inhibit the synthesis of negative-strand RNA more potently than that of positive-strand RNA and to change the component ratios of sgRNAs generated by both canonical and noncanonical template-switching events (Zhao et al., 2021b). The dominant effect of remdesivir on the synthesis of sgRNA3 may be derived from dysregulation in both canonical and noncanonical template-switching events in the replicon cells.

EIDD-2801, a prodrug of β-DN 4-hydroxycytidine (NHC), effectively suppressed viral replication in the transient and stable replicon assays (Fig. 3) and has been shown to potently inhibit SARS-CoV-2 replication *in vivo* and *in vitro* (Sheahan et al., 2020). The prodrug is intracellularly converted to the triphosphate form, which is a competitive substrate for CTP or UTP during the RdRp reaction. Thus, the metabolic pathway may be defective in 293T cells (Fig. 3A). On the other hand, EIDD-2801 dose-dependently inhibited SARS-CoV-2 particle production in VeroE6 cells expressing TMPRSS2 (Szemiel et al., 2021), suggesting that the VeroE6/Rep cell line is capable of supporting the metabolic activation of EIDD-2801.

In this study, we established transient and stable replicon systems of SARS-CoV-2. The transient replicon system will be available for the comparison of drug efficiency or replication activity among various cell types, while the stable replicon system using VeroE6/Rep3 cells is suggested to be suitable for applications with high-throughput screening. Our replicon systems will be helpful to advance knowledge in the field of research on the mechanism of SARS-CoV-2 replication and the development of effective antiviral agents or vaccines for clinical use.

## Acknowledgments

We thank M. Mori and M. Yoda for their assistance. We are also grateful to Jun-ichi Sakuragi, Kanagawa Prefectural Institute of Public Health, for providing SARS-CoV-2. This work was supported by the Japan Agency for Medical Research and Development (AMED) (20fk0108263, 20fk 0108518, 20fk0108471) and JST Moonshot R&D (JPMJMS2025).

## Author contributions

T.T., A.S. and T.S. performed the cloning and construction of vectors; A.S., Y.M., T.S. and K.T. provided critical materials and scientific insights; T.T. performed experiments; T. T., T.O., K.T. and K.M. analyzed the data; and T.T., K.T. and K.M. wrote the manuscript.

## STAR methods

### Lead Contact

Further information and requests for resources and reagents should be directed to and will be fulfilled by either of the Lead Contact, Kohji Moriishi.

### Materials Availability

All unique materials generated in this study are available from either of the lead contact with a completed Materials Transfer Agreement.

### Data Code and Availability

This study did not generate any unique datasets or code.

Experimental Model and Subject Details

Reagents and cells.

We purchased favipiravir, camostat mesylate and remdesivir from Cayman Chemicals (Ann Arbor, MI), EIDD-2801 (molnupiravir) from MedChemExpress (Monmouth Junction, NJ), recombinant human IFN-β from PeproTech (rocky Hill, NJ), and diamidino-2-phenylindole (DAPI) from Dojin Chemical (Kumamoto, Japan). The following antibodies were used in this study: mouse anti-SARS-CoV-2 nucleocapsid (N) monoclonal antibody (3A9; Cell Engineering Co. Osaka, Japan), mouse anti-SARS-CoV-2 nsp8 antibody (5A10; GeneTex, Irvine, CA), mouse anti-β-actin antibody (AC-74; Sigma–Aldrich), HRP-conjugated rabbit anti-mouse IgG antibody (Santa Cruz Biotechnology, Santa Cruz, CA) and Alexa 488-conjugated goat anti-mouse IgG antibody (Thermo Fisher Scientific, Waltham, MA). The human embryonal kidney 293T, human hepatoma Huh7 and African green monkey kidney VeroE6 cell lines were cultured in DMEM (Sigma-Aldrich, St. Louis, MO) supplemented with 10% FBS (Gibco, Cambrex, MD) and 1% sodium pyruvate (Sigma–Aldrich), 1% nonessential amino acid (Sigma–Aldrich) and 1% penicillin–streptomycin solutions (Nacalai Tesque, Kyoto, Japan). G418 (Nacalai Tesque) was further added to the culture medium at a final concentration of 1 mg/ml for the establishment and maintenance of the VeroE6/Rep cell line. All cell lines were cultured at 37°C in a 5% CO_2_ humidified atmosphere. SARS-CoV-2 strain Hu/DP/Kng/19-020 (GenBank accession no. LC528232.2) was kindly provided by Dr Jun-ichi Sakuragi at the Kanagawa Prefectural Institute of Public Health.

### Method details

#### Cloning, preparation and transfection of the BAC vector

Overviews of the design and construction of the replicon-BAC vector (pBAC-SCoV2-Rep-Reo) are shown in Figs. 1 and S1. The region including the viral replicase complex was amplified from the cells infected with SARS-CoV-2/Hu/DP/Kng/19-020 (GenBank accession no. LC528232.2). CPER enables direct synthesis of infectious cDNA of flaviviruses and SARS-CoV-2 from DNA fragments without *in vitro* ligation (Edmonds et al., 2013; Kotaki et al., 2021; Quan and Tian, 2009; Torii et al., 2021). First, we generated a pBAC cassette vector that encodes the cytomegalovirus (CMV) immediate-early promoter (CMVp) immediately upstream of the 5’ UTR of SARS-CoV-2 and the N gene followed by the 3’ UTR, synthetic 25-nt poly(A), the hepatitis delta virus ribozyme and BGH polyadenylation sequences, as the PCR template for fragment F10 (Fig. 1A), according to the method of Edmonds et al. (Edmonds et al., 2013). Eight fragments covering the replicase complex-encoding region were subcloned into pCR-blunt (Invitrogen) as templates for fragments F1 to F8 (Fig. 1A). Fragment F9, which is composed of, in order, a spacer (AceI site), the 3’-end portion of ORF8, TRS-B, the *Reo* gene, a spacer (BamHI site), the 3’-end portion of ORF8, and TRS-B, was subcloned into pCR-blunt (Fig. 1A, S1A). Eight fragments covering the region encoding the replicase complex (F1-to-F8 in Fig. 1A) were amplified by PCR using the primers listed in Table 1 and then cloned into pCR-Blunt (Invitrogen). To construct the pBAC cassette, the CMVp and the 5’-terminal region of SARS-CoV-2 (nt 1-683) were amplified by PCR from pcDNA3.1 and viral cDNA, respectively, combined by overlap PCR, and then cloned into pSMART-BAC v2.0 (Lucigen Corp., Middletown, WI). Subsequently, the 3’-terminal region of SARS-CoV-2 (nt 28,134-29,902) and polyadenylation sequence were amplified by PCR from the viral cDNA and pcDNA3.1, respectively, connected across the synthesized sequence consisting of 25-nt poly(A) and the hepatitis delta virus ribozyme (Almazan et al., 2006) by overlap PCR, and cloned into the BAC described above. The resulting pBAC cassette was used as the template for F10 production (Fig. 1A). Finally, the ten PCR fragments (F1 to F10) were mixed at an equal amount (0.05 pmol each) in 50 μl of the reaction buffer for PrimeSTAR GXL DNA polymerase (Takara Bio, Siga, Japan) and assembled *in vitro* by CPER (Edmonds et al., 2013; Torii et al., 2021). The CPER was carried out as follows: one cycle of 98°C for 2 min; 20 cycles of 98°C for 10 sec, 55°C for 15 sec and 68°C for 30 min; and one cycle of 68°C for 30 min followed by holding at 4°C. The resulting solution was extracted with phenol–chloroform–isoamyl alcohol, recovered by ethanol precipitation, dissolved in TE buffer, and introduced into BAC-Optimized Replicator v2.0 electrocompetent cells (Lucigen Corp.) according to the manufacturer’s protocol. The resulting vector was designated pBAC-SCoV2-Rep-Reo. DNA preparation was carried out using NucleoSpin (Takara Bio) and NucleoBond Xtra BAC Kits (Takara Bio) for small-scale and large-scale replicon-BAC vector preparation, respectively. Transfection of the replicon-BAC vector was performed using TransIT-LT1 reagent (Mirus, Madison, WI).

#### RT–PCR and quantitative RT–PCR (qRT–PCR)

Total RNA was extracted from cells using an RNeasy Mini Kit (Qiagen, Valencia, CA) and reverse transcribed using ReverTra Ace qPCR RT Master Mix (Toyobo, Tokyo, Japan). RT-PCR was performed to detect sgRNA3 and GAPDH mRNA by using Quick Taq HS Dye Mix (Toyobo). Quantitative RT–PCR was performed by using Fast SYBR Green Master Mix (Applied Biosystems, Foster City, CA) and a StepOne Plus Real-Time PCR System (Applied BioSystems). The expression level of sgRNA3 was calculated by the relative threshold cycle (ΔΔC_T_) method using GAPDH mRNA as an internal control.

The primers for RT–PCR and qRT–PCR are listed in Table 1.

#### Northern blotting

Total RNA was extracted from cells using TRIzol reagent (Invitrogen, Carlsbad, CA), precipitated, dissolved in RNase-free water and treated with a TURBO DNA-Free Kit (Thermo Fisher Scientific) according to the manufacturer’s protocol. The resulting preparations were then mixed with a formaldehyde loading dye (Ambion). The samples were denatured at 65°C for 15 min, chilled on ice, and electrophoresed through a 1% agarose gel containing formaldehyde. The separated RNA in the gel was stained with ethidium bromide (EtBr). After capillary transfer onto a positively charged nylon membrane (Roche, Mannheim, Germany), the membrane was subjected to UV fixation and then Northern blotting as reported previously (Otoguro et al., 2021). The signals were visualized with CDP-star (Roche) and LAS-4000 Mini (GE Healthcare, Tokyo, Japan). DIG-labeled RNA probes targeting the SARS-CoV-2 N gene were synthesized using a DIG RNA Labeling Kit (Roche). The N region was amplified with the primers listed in Table 1 and cloned into pBluescriptII. The plasmid was linearized with XhoI and used as the template for probe synthesis.

#### Statistical analysis

Quantitative experiments were carried out in triplicate and are indicated as the means ± standard deviations (SDs). All experiments were repeated at least three times, and the representative data of the independent experiments are shown. Statistical significance between two groups was determined by Student’s *t* test. Statistical significance for multiple comparisons was determined by one-way ANOVA with Dunnett’s post hoc test. A *p* value less than 0.05 was considered statistically significant (*, *p* < 0.05; **, *p* < 0.01; ns, no significance).

#### FACS analysis

The replicon cells or parent cells were harvested by trypsinization, washed with PBS and fixed with 4% paraformaldehyde. The cells were incubated with a blocking buffer (PBS with 5% FBS and 0.1% saponin) at room temperature for 60 min. The cells were then incubated with a dilution buffer (PBS with 1% BSA and 0.1% saponin) containing anti-SARS-CoV-2 N antibody (1:500) at room temperature for 90 min. After washing with PBS, cells were incubated with a dilution buffer containing Alexa 488-conjugated secondary antibody (1:500) at room temperature for 60 min and washed with PBS. The cells were analyzed with BD FACSCalibur and CellQuest software (BD Biosciences).

#### Others

Luciferase activity was measured using a *Renilla* luciferase assay system (Promega, Madison, WI) and a Luminescencer-Octa (ATTO, Tokyo, Japan) according to the manufacturers’ protocols. Cell viability was evaluated by an MTS assay as reported previously (Tanaka et al., 2021). The lysates of harvested cells for each experiment were prepared on ice with lysis buffer (25 mM Tris-HCl (pH 7.5), 150 mM NaCl, 1 mM EDTA, 0.1% SDS, 0.5% sodium deoxycholate, 1% NP-40 and 5% glycerol) supplemented with Complete protease inhibitor cocktail (Roche, Indianapolis, IN) and were subjected to Western blotting according to a previously described method (Otoguro et al., 2021). The replicon cells or parent cells were subjected to immunostaining after fixation according to a previously reported method (Otoguro et al., 2021).

## The supplemental information includes 2 figures

**Figure S1.**
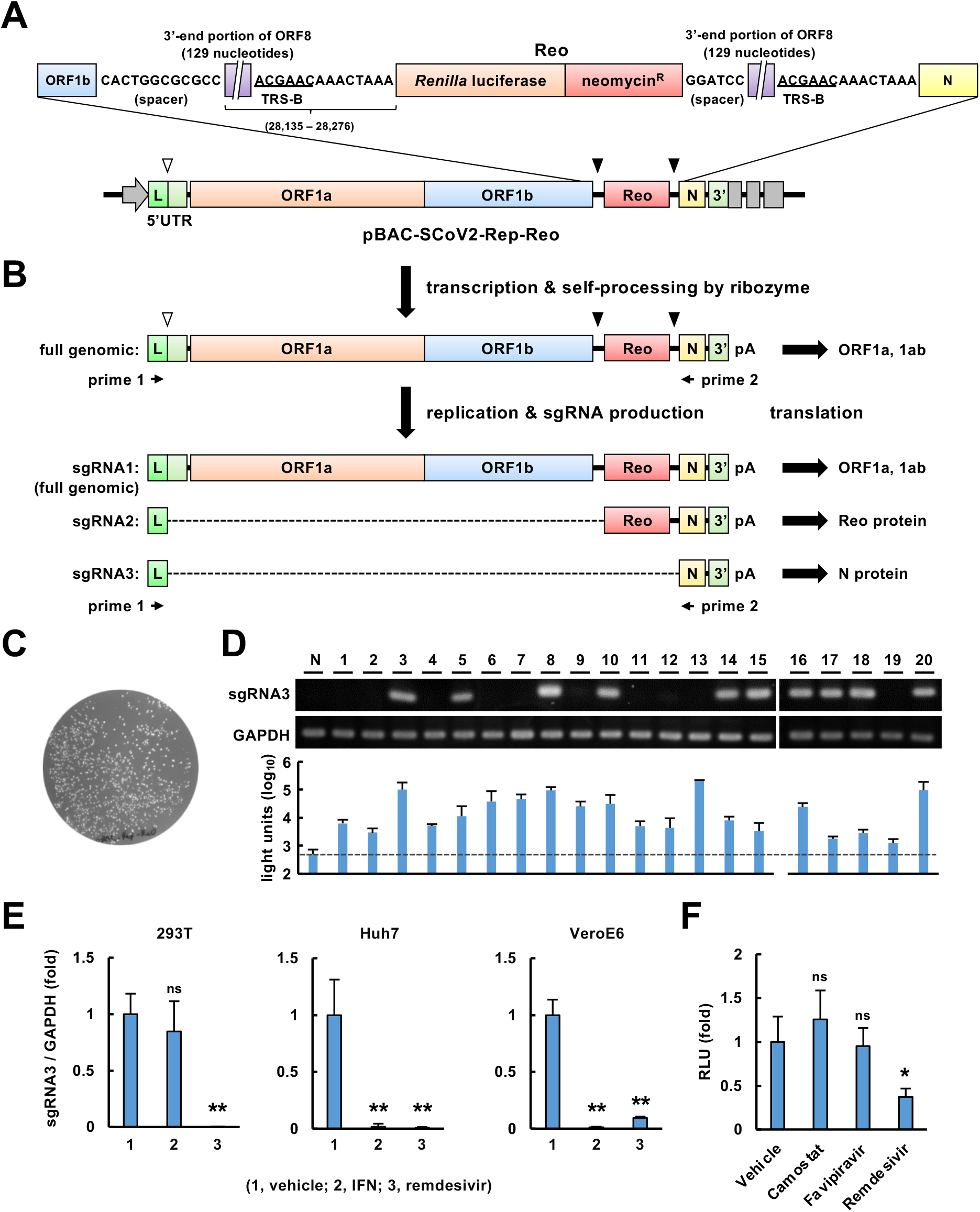
Construction, cloning and transient expression of pBAC-SCoV2-Rep-Reo. (A) Structure of the replicon-BAC vector. The region between the ORF1b and N genes in pBAC-SCoV2-Rep-Reo (the lower diagram) is magnified and shown in the upper diagram, corresponding to PCR fragment F9 (Fig. 1A). The numbers inside the parentheses represent the nucleotide positions in the SARS-CoV-2/Hu/DP/Kng/19-020 genome. (B) A schematic diagram of the production of sgRNAs. The upper diagram indicates full replicon RNA. The RNA transcribed under the control of the CMV promoter is processed by the ribozyme into full replicon RNA (sgRNA1). White and black arrowheads indicate the positions of TRS-L and TRS-B, respectively. sgRNA1 corresponding to full-length RNA and two sgRNAs (sgRNA2 and 3) are generated by discontinuous transcription mediated by the RNA-RNA interaction between TRS-L and TRS-B. sgRNA1, 2 or 3 serves as the translation template for polyprotein, Reo or N protein, respectively. Arrows indicate the location and orientation of the primers used for the detection and quantification of sgRNA3 (Table 1). (C) Direct transformation by the CPER product including pBAC-SCoV2-Rep-Reo. (D) Identification of *E. coli* transformants harboring pBAC-SCoV2-Rep-Reo. DNA preparations were isolated from twenty colonies shown in Fig. S1C and introduced into 293T cells. The transfected cells were harvested at 24 h post-transfection. These RNA preparations were subjected to RT–PCR using sgRNA3-specific primers 1 and 2 (upper) and to luciferase assays (lower). GAPDH mRNA was detected by RT–PCR as an internal control. The upper number and NC indicate the clone number and a mock-transfected negative control, respectively. (E) Evaluation of sgRNA3 expression. The relative level of sgRNA3 normalized to that of GAPDH mRNA was quantified by qRT–PCR. The indicated cell lines were transfected with the replicon-BAC vector and incubated with 0.1% DMSO (1), 100 U/ml IFN-β (2) or remdesivir (3) for 24 h. Remdesivir was applied to 293T cells at 0.5 μM and to Huh7 and VeroE6 cells at 5 μM. (F) Evaluation of luciferase activity. The replicon-BAC vector was transfected into 293T cells. The resulting cells were treated with 0.1% DMSO, 10 μM camostat mesylate, 10 μM favipiravir or 0.5 μM remdesivir at 6 h post-transfection and harvested at 30 h post-transfection. (E, F) Statistical significance was determined by one-way ANOVA with Dunnett’s post hoc test and comparison to the mock group (*, p < 0.05; **, p < 0.01; ns, no significance).

**Figure S2.**
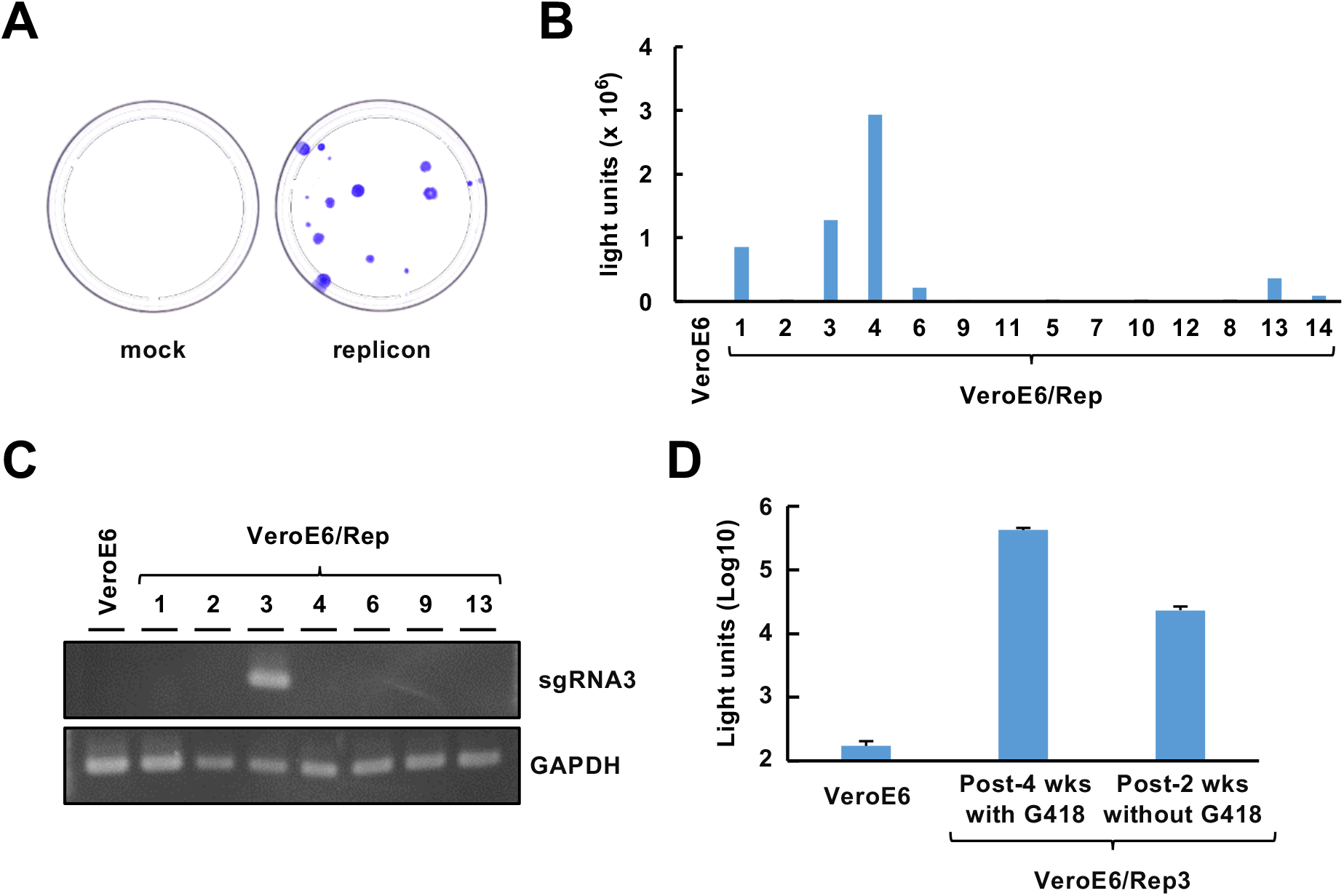
Screening for a VeroE6 cell line stably expressing the replicon. (A) G418-resistant colony formation by VeroE6 cells following transfection with the replicon vector (right side) or mock vector (left side). The cells were fixed with methanol and stained with crystal violet after continuous G418 exposure at 1 mg/ml. (B) Luciferase activities of parental VeroE6 cells and VeroE6-derived clones that were established from G418-resistant colonies following transfection with the replicon vector. The numbers below the columns represent the cell clone numbers. (C) Detection of sgRNA3 expression induction. Total RNA was extracted from VeroE6 cells or VeroE6-derived cell lines and subjected to RT–PCR analyses targeting sgRNA3 and GAPDH mRNA. GAPDH mRNA was detected as an internal control. (D) The stability of reporter expression in VeroE6/Rep3 cells. The cells were serially passaged during the indicated period in the presence or absence of 1 mg/ml G418. The luciferase activities of the cells were measured. The lysate of VeroE6 cells was subjected to luciferase activity assay as a background.

## References

Almazan, F., Dediego, M.L., Galan, C., Escors, D., Alvarez, E., Ortego, J., Sola, I., Zuniga, S., Alonso, S., Moreno, J.L., et al. (2006). Construction of a severe acute respiratory syndrome coronavirus infectious cDNA clone and a replicon to study coronavirus RNA synthesis. J Virol 80, 10900–10906.

Bakowski, M.A., Beutler, N., Wolff, K.C., Kirkpatrick, M.G., Chen, E., Nguyen, T.H., Riva, L., Shaabani, N., Parren, M., Ricketts, J., et al. (2021). Drug repurposing screens identify chemical entities for the development of COVID-19 interventions. Nat Commun 12, 3309.

Cao, X. (2020). COVID-19: immunopathology and its implications for therapy. Nat Rev Immunol 20, 269–270.

Cheng, Y.W., Chao, T.L., Li, C.L., Chiu, M.F., Kao, H.C., Wang, S.H., Pang, Y.H., Lin, C.H., Tsai, Y.M., Lee, W.H., et al. (2020). Furin Inhibitors Block SARS-CoV-2 Spike Protein Cleavage to Suppress Virus Production and Cytopathic Effects. Cell Rep 33, 108254.

Choy, K.T., Wong, A.Y., Kaewpreedee, P., Sia, S.F., Chen, D., Hui, K.P.Y., Chu, D.K.W., Chan, M.C.W., Cheung, P.P., Huang, X., et al. (2020). Remdesivir, lopinavir, emetine, and homoharringtonine inhibit SARS-CoV-2 replication in vitro. Antiviral Res 178, 104786.

Collier, D.A., De Marco, A., Ferreira, I., Meng, B., Datir, R.P., Walls, A.C., Kemp, S.A., Bassi, J., Pinto, D., Silacci-Fregni, C., et al. (2021). Sensitivity of SARS-CoV-2 B.1.1.7 to mRNA vaccine-elicited antibodies. Nature 593, 136–141.

Edmonds, J., van Grinsven, E., Prow, N., Bosco-Lauth, A., Brault, A.C., Bowen, R.A., Hall, R.A., and Khromykh, A.A. (2013). A novel bacterium-free method for generation of flavivirus infectious DNA by circular polymerase extension reaction allows accurate recapitulation of viral heterogeneity. J Virol 87, 2367–2372.

Gordon, C.J., Tchesnokov, E.P., Woolner, E., Perry, J.K., Feng, J.Y., Porter, D.P., and Gotte, M. (2020). Remdesivir is a direct-acting antiviral that inhibits RNA-dependent RNA polymerase from severe acute respiratory syndrome coronavirus 2 with high potency. J Biol Chem 295, 6785–6797.

Grossoehme, N.E., Li, L., Keane, S.C., Liu, P., Dann, C.E., 3rd, Leibowitz, J.L., and Giedroc, D.P. (2009). Coronavirus N protein N-terminal domain (NTD) specifically binds the transcriptional regulatory sequence (TRS) and melts TRS-cTRS RNA duplexes. J Mol Biol 394, 544–557.

Group, R.C. (2020). Lopinavir-ritonavir in patients admitted to hospital with COVID-19 (RECOVERY): a randomised, controlled, open-label, platform trial. Lancet 396, 1345–1352.

Harvey, W.T., Carabelli, A.M., Jackson, B., Gupta, R.K., Thomson, E.C., Harrison, E.M., Ludden, C., Reeve, R., Rambaut, A., Consortium, C.-G.U., et al. (2021). SARS-CoV-2 variants, spike mutations and immune escape. Nat Rev Microbiol 19, 409–424.

He, X., Quan, S., Xu, M., Rodriguez, S., Goh, S.L., Wei, J., Fridman, A., Koeplinger, K.A., Carroll, S.S., Grobler, J.A., et al. (2021). Generation of SARS-CoV-2 reporter replicon for high-throughput antiviral screening and testing. Proc Natl Acad Sci U S A 118.

Hoffmann, M., Hofmann-Winkler, H., Smith, J.C., Kruger, N., Arora, P., Sorensen, L.K., Sogaard, O.S., Hasselstrom, J.B., Winkler, M., Hempel, T., et al. (2021). Camostat mesylate inhibits SARS-CoV-2 activation by TMPRSS2-related proteases and its metabolite GBPA exerts antiviral activity. EBioMedicine 65, 103255.

Hoffmann, M., Kleine-Weber, H., Schroeder, S., Kruger, N., Herrler, T., Erichsen, S., Schiergens, T.S., Herrler, G., Wu, N.H., Nitsche, A., et al. (2020). SARS-CoV-2 Cell Entry Depends on ACE2 and TMPRSS2 and Is Blocked by a Clinically Proven Protease Inhibitor. Cell 181, 271–280 e278.

Huang, C., Wang, Y., Li, X., Ren, L., Zhao, J., Hu, Y., Zhang, L., Fan, G., Xu, J., Gu, X., et al. (2020). Clinical features of patients infected with 2019 novel coronavirus in Wuhan, China. Lancet 395, 497–506.

Kaptein, S.J.F., Jacobs, S., Langendries, L., Seldeslachts, L., Ter Horst, S., Liesenborghs, L., Hens, B., Vergote, V., Heylen, E., Barthelemy, K., et al. (2020). Favipiravir at high doses has potent antiviral activity in SARS-CoV-2-infected hamsters, whereas hydroxychloroquine lacks activity. Proc Natl Acad Sci U S A 117, 26955–26965.

Kokic, G., Hillen, H.S., Tegunov, D., Dienemann, C., Seitz, F., Schmitzova, J., Farnung, L., Siewert, A., Hobartner, C., and Cramer, P. (2021). Mechanism of SARS-CoV-2 polymerase stalling by remdesivir. Nat Commun 12, 279.

Kotaki, T., Xie, X., Shi, P.Y., and Kameoka, M. (2021). A PCR amplicon-based SARS-CoV-2 replicon for antiviral evaluation. Sci Rep 11, 2229.

Lei, X., Dong, X., Ma, R., Wang, W., Xiao, X., Tian, Z., Wang, C., Wang, Y., Li, L., Ren, L., et al. (2020). Activation and evasion of type I interferon responses by SARS-CoV-2. Nat Commun 11, 3810.

Martinez, M.A. (2020). Clinical Trials of Repurposed Antivirals for SARS-CoV-2. Antimicrob Agents Chemother 64.

McBride, R., van Zyl, M., and Fielding, B.C. (2014). The coronavirus nucleocapsid is a multifunctional protein. Viruses 6, 2991–3018.

Nguyen, H.T., Falzarano, D., Gerdts, V., and Liu, Q. (2021). Construction of a Noninfectious SARS-CoV-2 Replicon for Antiviral-Drug Testing and Gene Function Studies. J Virol 95, e0068721.

Osada, N., Kohara, A., Yamaji, T., Hirayama, N., Kasai, F., Sekizuka, T., Kuroda, M., and Hanada, K. (2014). The genome landscape of the african green monkey kidney-derived vero cell line. DNA Res 21, 673–683.

Otoguro, T., Tanaka, T., Kasai, H., Kobayashi, N., Yamashita, A., Fukuhara, T., Ryo, A., Fukai, M., Taketomi, A., Matsuura, Y., et al. (2021). Establishment of a Cell Culture Model Permissive for Infection by Hepatitis B and C Viruses. Hepatol Commun 5, 634–649.

Painter, W.P., Holman, W., Bush, J.A., Almazedi, F., Malik, H., Eraut, N., Morin, M.J., Szewczyk, L.J., and Painter, G.R. (2021). Human Safety, Tolerability, and Pharmacokinetics of Molnupiravir, a Novel Broad-Spectrum Oral Antiviral Agent with Activity Against SARS-CoV-2. Antimicrob Agents Chemother.

Pizzorno, A., Padey, B., Dubois, J., Julien, T., Traversier, A., Duliere, V., Brun, P., Lina, B., Rosa-Calatrava, M., and Terrier, O. (2020). In vitro evaluation of antiviral activity of single and combined repurposable drugs against SARS-CoV-2. Antiviral Res 181, 104878.

Quan, J., and Tian, J. (2009). Circular polymerase extension cloning of complex gene libraries and pathways. PLoS One 4, e6441.

Shannon, A., Selisko, B., Le, N.T., Huchting, J., Touret, F., Piorkowski, G., Fattorini, V., Ferron, F., Decroly, E., Meier, C., et al. (2020). Rapid incorporation of Favipiravir by the fast and permissive viral RNA polymerase complex results in SARS-CoV-2 lethal mutagenesis. Nat Commun 11, 4682.

Sheahan, T.P., Sims, A.C., Zhou, S., Graham, R.L., Pruijssers, A.J., Agostini, M.L., Leist, S.R., Schafer, A., Dinnon, K.H., 3rd, Stevens, L.J., et al. (2020). An orally bioavailable broad-spectrum antiviral inhibits SARS-CoV-2 in human airway epithelial cell cultures and multiple coronaviruses in mice. Sci Transl Med 12.

Shen, Z., Zhang, G., Yang, Y., Li, M., Yang, S., and Peng, G. (2021). Lysine 164 is critical for SARS-CoV-2 Nsp1 inhibition of host gene expression. J Gen Virol 102.

Siegel, D., Hui, H.C., Doerffler, E., Clarke, M.O., Chun, K., Zhang, L., Neville, S., Carra, E., Lew, W., Ross, B., et al. (2017). Discovery and Synthesis of a Phosphoramidate Prodrug of a Pyrrolo[2,1-f][triazin-4-amino] Adenine C-Nucleoside (GS-5734) for the Treatment of Ebola and Emerging Viruses. J Med Chem 60, 1648–1661.

Snijder, E.J., Decroly, E., and Ziebuhr, J. (2016). The Nonstructural Proteins Directing Coronavirus RNA Synthesis and Processing. Adv Virus Res 96, 59–126.

Szemiel, A.M., Merits, A., Orton, R.J., MacLean, O.A., Pinto, R.M., Wickenhagen, A., Lieber, G., Turnbull, M.L., Wang, S., Furnon, W., et al. (2021). In vitro selection of Remdesivir resistance suggests evolutionary predictability of SARS-CoV-2. PLoS Pathog 17, e1009929.

Tanaka, T., Okuyama-Dobashi, K., Motohashi, R., Yokoe, H., Takahashi, K., Wiriyasermkul, P., Kasai, H., Yamashita, A., Maekawa, S., Enomoto, N., et al. (2021). Inhibitory effect of a novel thiazolidinedione derivative on hepatitis B virus entry. Antiviral Res 194, 105165.

Thoms, M., Buschauer, R., Ameismeier, M., Koepke, L., Denk, T., Hirschenberger, M., Kratzat, H., Hayn, M., Mackens-Kiani, T., Cheng, J., et al. (2020). Structural basis for translational shutdown and immune evasion by the Nsp1 protein of SARS-CoV-2. Science 369, 1249–1255.

Torii, S., Ono, C., Suzuki, R., Morioka, Y., Anzai, I., Fauzyah, Y., Maeda, Y., Kamitani, W., Fukuhara, T., and Matsuura, Y. (2021). Establishment of a reverse genetics system for SARS-CoV-2 using circular polymerase extension reaction. Cell Rep 35, 109014.

Wahl, A., Gralinski, L.E., Johnson, C.E., Yao, W., Kovarova, M., Dinnon, K.H., 3rd, Liu, H., Madden, V.J., Krzystek, H.M., De, C., et al. (2021). SARS-CoV-2 infection is effectively treated and prevented by EIDD-2801. Nature 591, 451–457.

Yang, H., and Rao, Z. (2021). Structural biology of SARS-CoV-2 and implications for therapeutic development. Nat Rev Microbiol.

Yuan, S., Peng, L., Park, J.J., Hu, Y., Devarkar, S.C., Dong, M.B., Shen, Q., Wu, S., Chen, S., Lomakin, I.B., et al. (2020). Nonstructural Protein 1 of SARS-CoV-2 Is a Potent Pathogenicity Factor Redirecting Host Protein Synthesis Machinery toward Viral RNA. Mol Cell 80, 1055–1066 e1056.

Zhang, Y., Song, W., Chen, S., Yuan, Z., and Yi, Z. (2021). A bacterial artificial chromosome (BAC)-vectored noninfectious replicon of SARS-CoV-2. Antiviral Res 185, 104974.

Zhao, J., Guo, S., Yi, D., Li, Q., Ma, L., Zhang, Y., Wang, J., Li, X., Guo, F., Lin, R., et al. (2021a). A cell-based assay to discover inhibitors of SARS-CoV-2 RNA dependent RNA polymerase. Antiviral Res 190, 105078.

Zhao, Y., Sun, J., Li, Y., Li, Z., Xie, Y., Feng, R., Zhao, J., and Hu, Y. (2021b). The strand-biased transcription of SARS-CoV-2 and unbalanced inhibition by remdesivir. iScience 24, 102857.

Zuniga, S., Cruz, J.L., Sola, I., Mateos-Gomez, P.A., Palacio, L., and Enjuanes, L. (2010). Coronavirus nucleocapsid protein facilitates template switching and is required for efficient transcription. J Virol 84, 2169–2175.

